# Habitat structural complexity increases age-class coexistence and population growth rate through relaxed cannibalism in a freshwater fish

**DOI:** 10.1101/2023.07.18.549540

**Authors:** Eric Edeline, Yoann Bennevault, David Rozen-Rechels

**Affiliations:** DECOD (Ecosystem Dynamics and Sustainability), INRAE, IFREMER, Institut Agro, Rennes, France; U3E, INRAE, Agrocampus Ouest, Rennes, France; Sorbonne Université, CNRS, IRD, INRAE, Université Paris Est Créteil, Université Paris Cité, Institute of Ecology and Environmental Sciences of Paris (iEES-Paris), Paris, France

**Keywords:** Anthropogenic change, Biodiversity, Body size, Cannibalism, Competition, Conservation, Density-dependent population dynamics, Habitat degradation

## Abstract

Structurally-complex habitats harbour more taxonomically-diverse and more productive communities, a phenomenon generally ascribed to habitat complexity relaxing the strength of inter-specific predation and competition. Here, we extend this classical, community-centred view by showing that positive complexity-diversity and complexity-productivity relationships may also emerge from between-age-class, *intra-*specific interactions at a single-population level. In the laboratory, we show that medaka fish (*Oryzias latipes*) are strongly cannibalistic in complexity-free habitats, and that cannibalism may occur over a wide range of victim/cannibal body size ratios. In replicated outdoor pond populations, we manipulated habitat structural complexity using floating artificial structures, which selectively hampered movements of large-bodied medaka. Habitat complexity relaxed the strength of cannibalism, resulting in (1) increased survival of age-0+ individuals, (2) elevated age-class diversity, (3) increased population growth rate, and (4) dampened negative density-dependence in the stock-recruitment relationship reflecting elevated habitat carrying capacity. The resultant higher population density in complex habitats was associated with increased competition for food among both age-0+ and age-1+ individuals. Our results highlight that positive complexity-diversity and complexity-productivity relationships may be considered as a generally-emergent property of size-structured populations and communities in which a larger body size brings a predation or interference advantage. Hence, enhancement of habitat structural complexity may be seen as a pivotal management strategy not only in favour of taxonomic diversity, but also to increase the productivity and resilience of exploited populations and to improve the conservation status of endangered species.

## INTRODUCTION

Anthropogenic changes and biodiversity loss are often congruent with a simplification in the physical structure of habitats. For instance, agriculture transforms complex, mature forests into pastures or grain fields that, eventually, may return to secondary forests which are structurally simpler than mature forests (Colorado Zuluaga and Rodewald 2015). In aquatic or terrestrial habitats, top-predator extirpation may have cascading effects resulting in a loss of the large autotrophic organisms that structure space (Estes et al. 2011). In oceans, eutrophication, acidification, bottom trawling and dredging all tend to destroy the complex structure provided by benthic habitats such as coral reefs (National Research Council 2002, Rogers et al. 2014). In freshwaters, channelisation for hydropower, shipping, urban development or log-driving tremendously simplifies river habitats, and eutrophication may drive a loss of submerged macrophytes which are key to structuring rivers, ponds or lakes (Scheffer 2004).

In parallel, many studies report that structurally-complex habitats favour increased species coexistence and higher numerical abundances and biomass in animal communities. This positive effect is generally ascribed to habitat complexity relaxing the strengths of both predation and competition, and thus allowing more species to coexist through increasing surface area, niche diversity and the spatial partitioning of limiting resources (Smith 1972, Crowder and Cooper 1982, Diehl 1988, 1992, Heck and Crowder 1991, Hixon and Menge 1991, Janssen et al. 2007, Kovalenko et al. 2012, Reichstein et al. 2013, Rogers et al. 2014, Soukup et al. 2022).

This is because highly-structured habitats make prey less detectable visually and hinder movements and encounters (Stenseth 1980, Bartholomew et al. 2000, St Pierre and Kovalenko 2014, Soukup et al. 2022), such that predators may be restricted to ambush hunting only (Schultz et al. 2009, Soukup et al. 2022). In particular, in complex habitats larger-bodied individuals disproportionately suffer from a reduced agility and from a reduced accessibility to microhabitats and crevices (Rogers et al. 2014, Soukup et al. 2022). Hence, structurally-complex habitats decrease predator attack rates, resulting in a reduced risk of prey overexploitation at all trophic levels and in relaxed apparent competition among prey, i.e., relaxed indirect competition through supporting a common predator (Holt 1987).

Historically, research on the ecological effects of habitat structural complexity has focused on interspecific interactions with more limited consideration for intraspecific interactions. This is despite that many populations are size-structured due to both overlapping age classes and due to heterogeneous somatic growth rates within age classes (Ebenman and Persson 1988). In such size-structured populations, apparent competition is by definition not possible, but large-bodied individuals often dominate in interference competition (Post et al. 1999, Le Bourlot et al. 2014), and may even cannibalize smaller-bodied conspecifics (Fox 1975, Smith and Reay 1991, Wise 2006). It is therefore likely that, in size-structured populations, just like in size-structured communities, a high habitat structural complexity impedes the dominance of large-bodied individuals.

Accordingly, a few studies suggest that habitat structural complexity increases age-class diversity in the form of enhanced juvenile-adult coexistence, supports larger population sizes, and favours population persistence in laboratory populations of the guppy *Poecilia reticulata* (Yamagishi 1976, Nilsson and Persson 2013) and of the mosquitofish *Gambusia affinis* (Benoît et al. 2000). However, given the paucity of studies dealing with habitat complexity in size-structured populations, the generality of positive complexity-diversity and complexity-productivity relationships at the population level remains questionable. Additionally, the respective contributions of competition and cannibalism to the emergence of complexity-productivity relationships at the population level remain largely unexplored.

In this paper, we test the prediction that habitat structural complexity increases age-class diversity and increases population productivity in the Japanese medaka fish (*Oryzias latipes*), and that the underlying mechanisms involve a relaxation in the strengths of both interference competition and cannibalism. Our approach was to first evaluate the potential for medaka express size-dependent cannibalism in the laboratory, and then to quantify the effects of structural complexity on medaka population dynamics in outdoor-pond experiments. Our results suggest that habitat complexity increases age-class diversity and population productivity, but point to cannibalism as the only underlying mechanism.

## MATERIALS AND METHODS

### Medaka fish

The medaka is an oviparous fish belonging to the group of Beloniformes, a sister group of Cyprinodontiformes which includes killifishes (Kinoshita et al. 2009). Medaka naturally inhabit slow-moving fresh- and brackish-waters of South-East Asia. Juveniles and adults have a highly-overlapping diet of zooplankton, small benthic invertebrates and filamentous algae (Terao 1985, Edeline et al. 2016). Due to its high thermal tolerance and ease of manipulation, the medaka can be used for parallel experiments in the laboratory, where generation time is 2-3 months under optimal light, food and temperature conditions, and in outdoor ponds, where perennial populations maintain themselves over years under natural conditions without any artificial feeding (Bouffet-Halle et al. 2021). For these reasons, the medaka is a good model species for studies in genetics, developmental biology, ecology and evolution (Kinoshita et al. 2009, Renneville et al. 2016, 2020, Diaz Pauli et al. 2019, 2020, Le Rouzic et al. 2020, Evangelista et al. 2020a, b, 2021, Bouffet-Halle et al. 2021).

### Cannibalistic behavioural assays and predation window

Trophic interactions are often size-dependent (Woodward et al. 2005), such that the prey/predator body-length ratio is constrained to a specific range called the ‘‘predation window” (Claessen et al. 2000). The lower and upper limits of the predation window have far-reaching consequences for population dynamics and individual life histories, respectively (Claessen et al. 2002). We performed cannibalistic behavioural assays to determine whether medaka are cannibalistic under no-complexity conditions and, if so, to estimate the limits of their predation window.

Cannibalistic assays were performed at the Centre de recherche en Ecologie Expérimentale et Prédictive (CEREEP, https://www.cereep.bio.ens.psl.eu/) using the Kiyosu medaka strain (Toyohashi, Aichi Prefecture, Japan, see Bouffet-Halle et al. 2021). Temperature was 21°C (±1.5 SD) and the light regime followed the natural photoperiod at the time of experiment (12-13 ^th^ March 2013), i.e., dark from 17:00 to 9:00. Adult medaka were measured individually for standard body length (Sdl, from the tip of the snout to the base of the caudal peduncle, n = 45 adults, mean ± SD = 22.8 mm ± 4.2), determined for sex according to their secondary sexual characters (Yamamoto 1975, Kinoshita et al. 2009), and were kept separately in a 0.5 L complexity-free aquarium filled with dechlorinated tap water. After 72h of adult fastening (i.e., at 15:30 on March 12^th^), two medaka newborns where chosen randomly from a pool of 2-10 days old larvae, were measured individually for Sdl (n = 90 larvae, mean ± SD = 5.6 mm ± 0.7), and were randomly-deposited with an adult in one of the 45 aquariums. It was not possible to follow larvae individually, and we therefore counted the number of surviving larvae in each aquarium after 2.5 hours, and then every 3 hours during 17.5 hours (yielding n = 270 observations). This approach made it possible to study the kinetics of larvae survival in each aquarium.

We used the relationship between larvae survival probability and victim/cannibal Sdl ratios (taking the average Sdl for the two larvae) to estimate the lower (δ) and higher (ε) limits of medaka predation window, as well as the optimal victim/cannibal Sdl ratio (ϕ), as defined by Claessen et al. (2002). Specifically, using the probability of a cannibalistic attack estimated at first census (2.5 hours of cannibalistic assays), i.e., when most of size-dependency in cannibalism was expressed (see Methods and Results), we arbitrarily defined δ and ε as the lower-and higher end Sdl ratios, respectively, at which cannibalistic probability becomes less than 0.05, and ϕ as the Sdl ratio at which cannibalistic probability is maximal.

### Geotextile to manipulate habitat complexity

During outdoor-pond experiments (see below), we manipulated habitat structural complexity using tiles of Maccaferri MacMat®, a high-porosity geotextile used in soil-erosion control. This floating geotextile forms a 3D surface with regularly-spaced, 12-13 mm-deep hills and valleys (Fig. 1A). The 3D surface is made from an irregular mesh of polyamide threads, which present a multiplicity of holes of various sizes through which fish can swim (Fig. 1A, Appendix 1). Such a floating artificial structure is well adapted to providing shelters to medaka, which is a surface-dwelling fish species (Yamamoto 1975, Kinoshita et al. 2009).

**Fig. 1.**
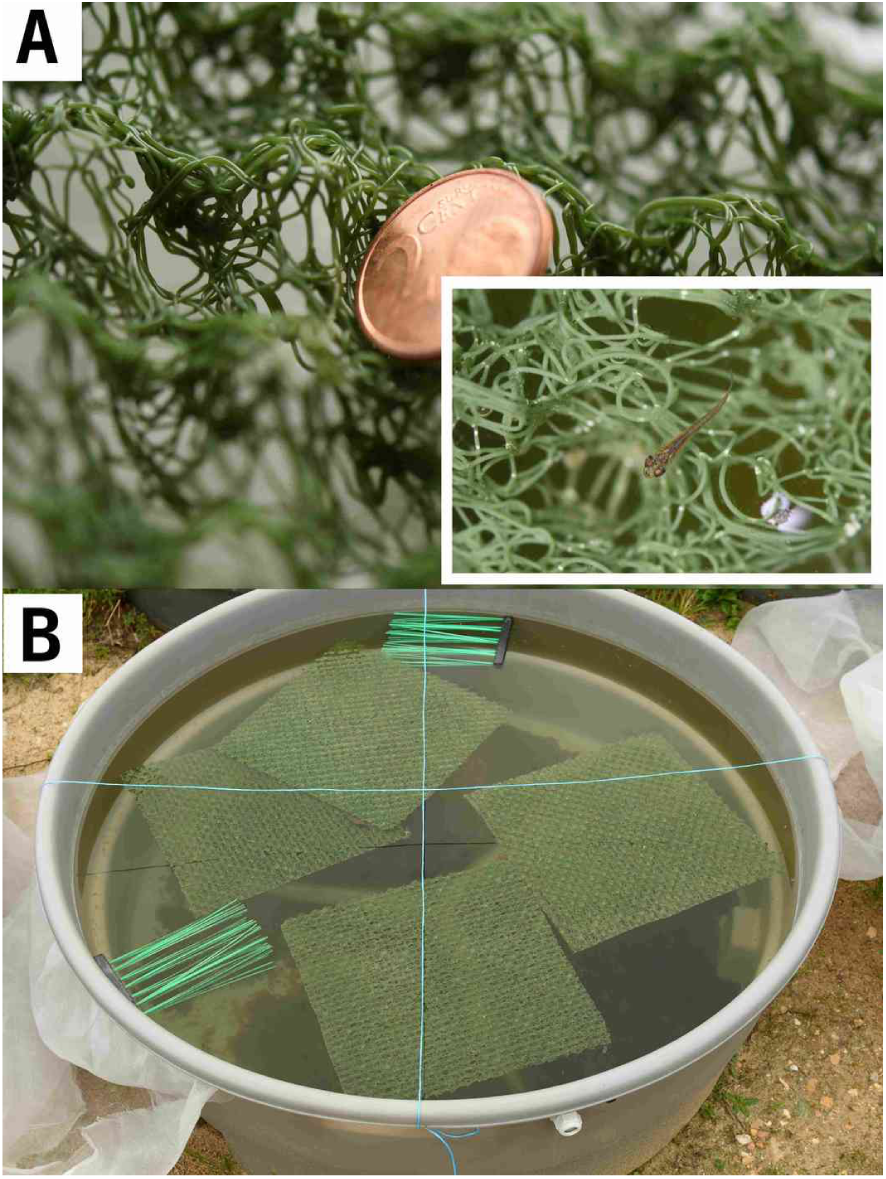
Geotextile manipulation of habitat structural complexity in pond medaka populations. **A:** Detailed structure of the geotextile used as artificial refuges to manipulate habitat complexity. The 2 euro-cent coin has a diameter of 22.25mm. The inset shows a juvenile medaka swimming through the geotextile. For a detailed analysis of geotextile structure, see Appendix 1. **B:** A high-complexity pond during the 2021 experiment, hosting single-layered tiles of geotextile, as well as two floating plastic brushes that ensured that spawning substrates for medaka were non-limiting. In 2022, high-complexity habitats were created using five-layered geotextile mats (see Appendix 1). NB: blue ropes ensured that the insect-proof net that covered ponds did not touch water surface.

A detailed comparison of geotextile structure with the structure provided by various types of natural vegetation is beyond the scope of this study. However, we note that near-surface or floating vegetation mats may be formed by riverbank-tree roots, accumulations of dead wood or other floating debris, by floating macro-algae (*Spirogyra* sp. or *Oedogonium* sp), or by macrophytes, be they ground-rooted (e.g., *Myriophylumm spp*.) or strictly-floating (e.g., *Utricularia vulgaris* or *Ceratophyllum demersum*) in natural water bodies.

To gain some insights into the mechanisms through which the geotextile could affect medaka behaviour, we measured the distribution of geotextile’s hole sizes (in 2D, as seen from a top view), and compared it with the distributions of medaka body width and height, which are the largest horizontal and vertical linear distances, respectively, perpendicular to the the fish normal direction of motion (Bartholomew et al. 2000). The distributions of medaka body width and height were fully included in geotextile’s hole-size distribution, suggesting that even the largest-bodied medaka can swim through the geotextile (Appendix 1). However, small-bodied medaka had many more holes available to swim through, suggesting that the geotextile would selectively hamper the movement of the largest medaka.

To test this hypothesis, we performed behavioural assays aiming at measuring the size-dependency of medaka ability to swim through a single layer of geotextile. The results confirmed that medaka of all sizes can swim through the geotextile, but that medaka larger than ca. 15 mm in Sdl need much more time to find their way through the geotextile (Appendix 1). Hence, the geotextile selectively hampered the movements of larger-bodied medaka, exactly as natural habitat structures selectively hamper the movement of large-bodied predators (Crowder and Cooper 1982, Diehl 1988, 1992, Heck and Crowder 1991, Bartholomew et al. 2000, Kovalenko et al. 2012, Rogers et al. 2014). Although we did not specifically test for visual effects, we hypothesize that the geotextile further interfered with medaka ability to see or sense small-bodied conspecifics (Bartholomew et al. 2000).

We quantified the effects of geotextile-manipulated habitat complexity on medaka population dynamics during two separate outdoor-pond experiments (16, 2 m^2^ circular ponds, see below). In 2021, we used single-layered geotextile tiles (12-13 mm deep) of varying surface to induce a complexity contrast. Specifically, single-layered tiles of geotextile covered either a low proportion of pond surface (from 3.1 to 9.9 %, mean 8.6 %, hereafter low-complexity treatment, n = 8 ponds) or a large proportion of pond surface (from 36.2 to 50 %, mean 39.4 %, n = 8 ponds, hereafter high-complexity treatment, Fig. 1B).

In 2022, to induce a more drastic complexity contrast, we used single-layered geotextile tiles in low-complexity habitats and five-layered geotextile mats (90-100 mm deep) in high-complexity habitats (Appendix 1). Specifically, the low-complexity treatment consisted of 0.063 m^2^ of single-layered geotextile tiles (3.1 % of pond surface), while the high-complexity treatment consisted of 0.750 m^2^ of five-layered geotextile tiles (amounting to 187.5 % of pond surface). Note that ponds were 500-600 mm deep (see below), such that most of the water column was unstructured in both the low- and high-complexity treatments.

In both 2021 and 2022, we added two floating plastic brushes to the ponds to avoid a limitation by the spawning substrate (Fig. 1B).

### Pond ecosystems and medaka populations

On 31^st^ March 2021, 12 circular, 2 m^2^ ponds (H = 0.60 m, diam. = 1.65 m, vol. = 1 m^3^) were installed at the U3E experimental research unit (https://www6.rennes.inrae.fr/u3e_eng/), filled with dechlorinated tap water, and seeded with 2 L of a mixture of benthic detritus (decaying leaves and phytoplankton), benthic organisms and plankton collected in mature, fishless ponds using a kick net (0.3 mm mesh). Benthos and plankton additions were repeated on April 2^nd^, 9^th^ and 13^th^. Additionally, on three occasions (April 2^nd^, 9^th^ and 15^th^) each pond received a solution of KH_2_PO_4_ amounting to 10 µg L^-1^ of phosphorus. Enrichment was complemented on April 13^th^, 19^th^ 27^th^ and June 4^th^ with additions of 1 L of an algal mixture (*Chlorella* sp. and *Desmodesmus* sp). All ponds were covered with an insect-proof net (1 mm mesh size). On July 1^st^ 2021, 4 more ponds were installed and submitted to the same seeding and enrichment treatments. Hence, the experiment included 16 ponds in total.

On March 31^st^ 2021 (July 8^th^ for the four extra ponds), from 18 to 60 adult medaka (mean = 37) born in 2020 (hereafter “age-1+”) were randomly introduced in ponds. Stocking numbers were n = 35 ± 12 fish at low complexity and n = 39 ± 14 fish at high complexity (estimate = 0.107, SE = 0.174, t-value = 0.617, p-value = 0.547). To vary the genetic background of the fish populations, we used medaka originating from the Kiyosu population, as well as five other strains corresponding to different locations in Japan and provided by the Japanese National Bioresource Project^1^: Hamochi, Higashidori, Inawashiro, Tokamachi, and Kushima. The six strains were not mixed in a pond, but seeded each in two separate ponds (one low-complexity pond and one high-complexity pond), except the Kiyosu strain that was seeded in 6 ponds (three low-complexity ponds and three high-complexity ponds). Preliminary analyses of the 2021 data showed that medaka strains had no significant effect on population growth rate or body sizes (Appendix 2), and we chose to discard the strain effect from our subsequent analyses and experimental design (see below).

From November 15^th^ to 17^th^ 2021, i.e., after the reproductive period, all fish from each pond were sampled using both hand nets and a seine net. To reduce fish handling and to provide fast and reproducible body-size measurements, fish were live-photographed in batches in a transparent tray set above a light source, and were released in their pond. These fish included a mixture of both age-1+ adults that were survivors from initial introductions and age-0+ recruits that were born in the ponds. Each photograph was analysed with ImageJ to automatically fit an ellipse to each fish shape using the “fit-ellipse” tool (Schneider et al. 2012). Sdl was equated to ellipse major distance.

On March 22^nd^ 2022, the 16 ponds were dried, and water and sediments were mixed and randomly redistributed among the 16 ponds. This was the start of a long-term experiment requiring identical initial conditions among ponds. Therefore, we homogeneously mixed medaka strains in each pond by seeding 3, 8, 7, 15, 8 and 25 fish from the Hamochi, Higashidori, Inawashiro, Tokamachi, Kushima and Kiyosu strains, respectively (66 fish per pond). Fish were batch-photographed for Sdl measurement using ImageJ on November 8^th^ and 9^th^ 2022, and were released in their pond to pursue the long-term experiment.

### Filamentous algae

Filamentous algae represent an important food source for medaka, both in the wild (Terao 1985, Edeline et al. 2016) and in mesocosms (Bouffet-Halle et al. 2021). In November 2022, before medaka fishing, two independent observers visually estimated the percentage of pond surface covered by filamentous algae in each pond.

### Statistical analyses

#### Cannibalistic behavioural assays

From cannibalistic assays in the laboratory, we estimated how the probability for an average-sized larvae to be cannibalized changed with (i) the victim-to-cannibal Sdl ratio, (ii) the time elapsed since the start of the assays, and (iii) the sex of the cannibal. To that aim, we used an overdispersed binomial (logit link) generalized additive model (GAM) of the form:

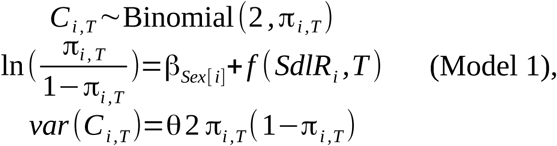

where ln is the natural logarithm, *C* is the number of larvae eaten, *i* indexes an aquarium (and cannibal individual), *T* indexes the time of census (*i*×*T* = 270 observations), π is cannibalism probability, β*_Sex_*_[_ *_i_*_]_ is a sex effect of cannibal individual *i*, *SdlR* is mean victim/cannibal Sdl ratio, *f* is a tensor product of natural cubic splines with 3 knots, which was parsimonious in terms of smoother wigliness and, at the same time, captured the essential features of the cannibalism-size relationship. Finally, θ is a positive parameter accounting for a slight overdispersion in the data. Estimating a different *f* for each sex increased model’s generalised cross-validation score, and we thus preferred the simpler model including *Sex* as a fixed effect on the intercept.

We fitted Model 1 using quasi-likelihood (quasibinomial family) in the mgcv library of the R software version 4.2.1 (Wood 2017, R Core Team 2024). This model explained 24.5 % of the deviance in the number of larvae eaten. The significance of the effects included in the model was assessed using a standard *t*-test (as provided by the summary function in R).

#### Pond populations

We analysed the effects of habitat structural complexity on pond medaka population dynamics in two steps. Step one aimed a gaining a general understanding of unstructured population dynamics using a classical Ricker model of total population numbers (which were measured with no error). As a second step, we tried to gain a more detailed understanding of the mechanisms behind population dynamics by deciphering age structure in the populations. We did so using a Gaussian mixture model that estimated individual age from body-size distributions.

At step one, we specifically explored unstructured medaka population dynamics using the Ricker logistic equation (Case 2000):

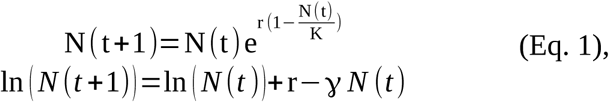

where *N* (*t* +1) is total fish number after reproduction (recruitment), *N* (*t*) is the number of age-1+ medaka parents initially introduced before reproduction (stock), r is maximum population growth rate, and γ=r/ K measures the strength of negative density-dependence in the population.

We tested for an effect of habitat structural complexity on the density-*in*dependent r parameter, and on the density-dependent γ parameter using a Poisson GLM:

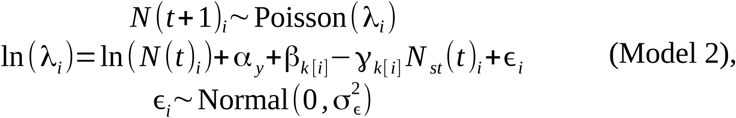

where *i* indexes pond-by-year combinations (n = 32), *y* indexes the year of experiment (2021 and 2022), α *_y_* and β*_k_*[*_i_*] are year-of-experiment and habitat-complexity effects on population growth rate, respectively, γ*_k_* [*_i_*] captures a habitat complexity-by-density interaction, and 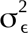 is an overdispersion parameter. To avoid any intercept-slope correlation, we standardized *N* (*t*) to zero mean (hence the *N_st_* (*t*) notation in Model 2 equation). As a consequence of this standardisation, the α and β parameters in Model 2 do not estimate the Ricker stock-recruitment relationship at origin, but at an average level of the stock.

As stressed above, the Ricker-logistic approach in Model 2 is unstructured by lumping together age-1+ parents and their age-0+ progeny in *N* (*t* +1), and is therefore unable to adult survival from the *per capita* production rate of age-0+ recruits (i.e., number of age-0+ at time *t*+1 / number of introduced age-1+ at time *t*). To gain a deeper understanding of medaka population dynamics, we inferred individual ages at *t*+1 from the information present in the body-size distributions. At this step two of our analysis, we deciphered mixtures of size distributions for age-1+ and age-0+ fish, and their response to habitat complexity, using a log-normal mixture model fitted to standard body lengths Sdl:

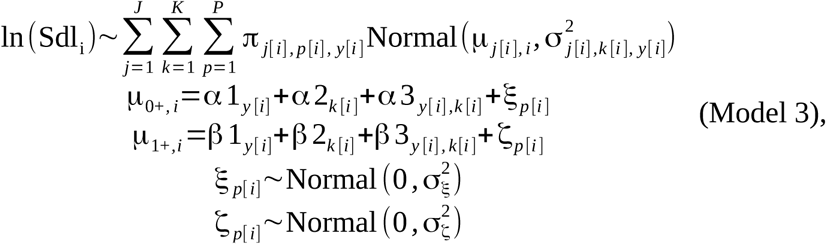

where *i* indexes individual fish (n = 3423), *j* indexes age groups (age 0+ *vs*. age 1+, *J* = 2), *k* indexes the complexity treatment (low vs. high, *K* = 2), and *p* indexes ponds (*P* = 16). π *_j_* _[_*_i_*_]_ *_, p_* _[_*_i_* _]_*_, y_*_[_ *_i_*_]_ is the proportion of age *j* fish in pond *p* and year of experiment *y* such that, for each *p* and 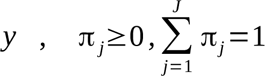. The α 1_y [i]_ and β1_y[i]_ parameters capture year-of-experiment effects (2021 vs. 2022) on mean body lengths at age-0+ and age-1+, respectively. The α 2_*k [i]*_ and β2_k[i]_ parameters capture the effects of habitat structural complexity on mean body lengths at age 0+ and age 1+, respectively. The α 3 *_y_*_[_*_i_* _]_*_, k_*_[_ *_i_*_]_ and β3 *_y_*_[_*_i_* _]_*_, k_*_[_ *_i_*_]_ parameters capture complexity-by-year interactions on mean body lengths at age 0+ and age 1+, respectively. The ξ*_p_* _[_*_i_* _]_ and ζ *_p_* _[_*_i_*_]_ parameters capture random pond effects on mean body lengths at age 0+ and age 1+, respectively. Finally, Model 3 included heteroscedasticity in the form of a triple age class-by-complexity-by-year interaction on residual variance σ^2^ (i.e., eight variances were estimated separately).

Estimates for the number of individuals in each age class in each pond and each year were computed from Bayesian posterior estimates (see below) of π *_j_* as 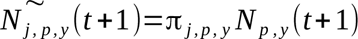, where *N_p, y_* is total fish number. To separate the effects of habitat complexity on *per capita* production rate of age-0+ recruits and on age-1+ parent survival, we used two separate Ricker-logistic models similar to Model 2. We modelled the median posterior number of age-0+ recruits 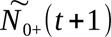 as:

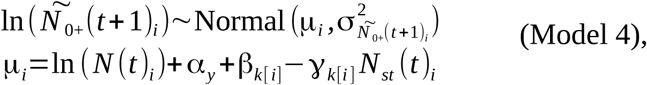

Where 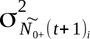 is the posterior variance in 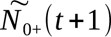, supplied here as data so as to propagate estimation uncertainty, α *_y_* and β*_k_*_[_ *_i_*_]_ capture the effects of year-of-experiment and habitat complexity, respectively, on *per capita* production rate of age-0+ recruits, evaluated at an average level of the parental stock (i.e., at *N_st_* (*t*)=0). The γ*_k_* _[_*_i_* _]_ captured the density-by-habitat complexity interaction on *per capita* production rate of age-0+ recruits.

Following a similar rationale, we modelled the median posterior number of surviving age-1+ fish after reproduction 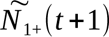 as:

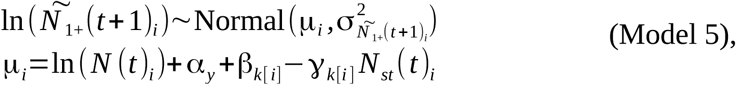

Where 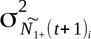 is posterior variance in 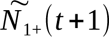, intercepts α and β capture the year-of-experiment and habitat-complexity effects on mean, ln-transformed survival probabilities of age-1+ fish through the reproductive period, and slopes γ_k [ i ]_ capture the habitat complexity-by-density interaction on age-1+ survival probabilities. We preferred Model 5 to a Binomial or a Beta models, which are more naturally adapted to modelling probabilities, but can not straightforwardly incorporate estimation uncertainty 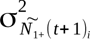 However, as a complement to Model 5, we provide a Binomial analysis of age-1+ survival probabilities in Appendix 3.

We fitted Models 2-5 using Markov Chain Monte Carlo (MCMC) in JAGS 4.3.0 (Plummer 2003) through the jagsUI package (Kellner 2019). Priors were chosen to be weakly informative, except in mixture Model 3, where we imposed the following constraints: (i) mean standard body length was smaller at age 0+ than at age 1+ (i.e., α_1_ <β_1_ and α_2_ <β_2_), (ii) body-length variance was larger at age 0+ than at age 1+, as evidenced by a visual inspection of size distributions (i.e., 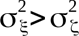), and (iii) the number of age-1+ individuals after reproduction at time *t*+1 was not larger than before reproduction at time *t* (i.e., 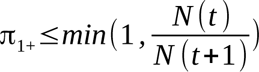). We further prevented label *N* (*t* +1) switching by assigning age-class 0+ to fish shorter than 10 mm and age-class 1+ to fish longer than 30 mm (Edeline et al. 2016, Bouffet-Halle et al. 2021). We ran 3 parallel MCMC chains until parameter convergence was reached, as assessed using the Gelman–Rubin statistic (Gelman and Rubin 1992).

We assessed goodness of fit of Models 2-5 by using a Bayesian P-value (Gelman et al. 1996). Briefly, we computed residuals for the actual data as well as for synthetic data simulated from estimated model parameters (i.e., residuals from fitting the model to ‘‘ideal’’ data). The Bayesian P-value is the proportion of simulations in which ideal residuals are larger than true residuals. If the model fits the data well, the Bayesian P-value is close to 0.5. Bayesian P values for Models 2-5 were 0.49, 0.52, 0.12 and 0.07, respectively, indicating fits ranging from excellent to fair only (Gelman et al. 1996). The lower fit of Models 4 and 5 is explained by the fact that error variance was not freely estimated but supplied as data in these models.

The significance of the effects included in Models 2-5 was assessed using MCMC p-values (not to be confounded with the Bayesian P-value above), which quantify posterior overlap with zero. Specifically, at each MCMC iteration and for each model parameter, we computed Δ as the difference between posterior parameters under low and high habitat complexity. We then computed MCMC p-values as twice the proportion of Δ for which the sign of Δ was opposite to that of its mean value.

Finally, the effect of habitat complexity on the percentage of pond surface covered by filamentous algae *Pr* was modelled using a Beta GLMM:

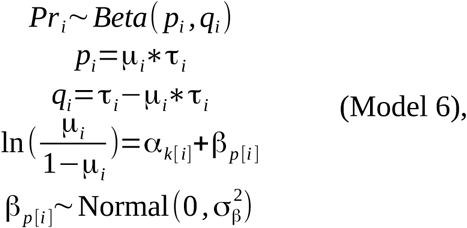

where *i* indexes an observer-by-pond combination (n = 32 observations), α*_k_*_[_ *_i_*_]_ captures the effect of habitat structural complexity (low vs. high), and β*_p_*_[_ *_i_*_]_ is a random pond effect (n = 16 ponds). We fitted Model 6 in the glmmTMB library of R (Brooks et al. 2017), and evaluated significance of the complexity effect using the summary function of R.

## RESULTS

### Cannibalistic behavioural assays and predation window

The relationship between medaka cannibalistic behaviour and the victim/cannibal standard body length (Sdl) ratio predicted by GAM Model 1 was nonlinear, and changed during the course of cannibalistic assays (Fig. 2A). At first census, after 2.5 hours of exposure, freely-adjusting splines showed that the relationship followed a bell-shaped curve, thus validating the hypotheses of the predation window (Fig. 2B).

**Fig. 2.**
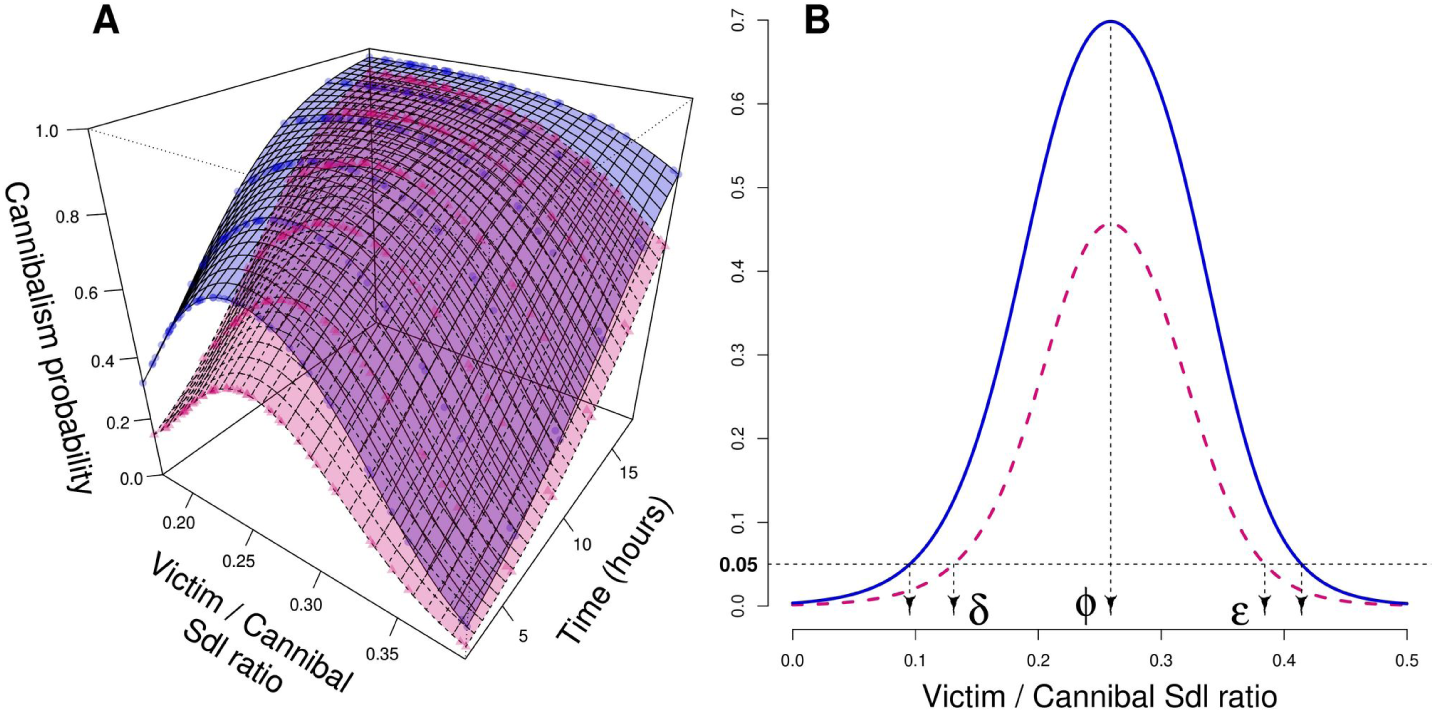
Cannibalistic behaviour in the laboratory with no habitat structure. **A:** Surface plot for the nonlinear interaction between victim/cannibal standard body length (Sdl) ratio and time of exposure on the probability for cannibalism to occur. Blue solid lines: male cannibals; Pink dashed lines: female cannibals. The surfaces were predicted from Model 1. Dots show the fitted values: Circles: male cannibals; Triangles: female cannibals. **B:** Defining parameters of the predation window from predicted cannibalism probability surfaces. The cannibalistic curves are the same as in Fig. 2A for the fist census (2.5 hours of cannibalistic assays). δ and ε are the lower and upper limits of the predation window, arbitrarily defined as the lower- and upper-end Sdl ratios, respectively, at which cannibalistic probability becomes less than 0.05 (horizontal dashed line). ϕ is the optimal victim-to-cannibal Sdl ratio. The resultant predation window is plotted in Fig. 3. Note that the higher cannibalistic voracity in males results in a wider predation window.

As time of victim to cannibal exposure was increasing, the relationship progressively became body-size independent. By the end of the assays (17.5 hours of exposure), victim survival probability was very low under almost all Sdl ratios (Fig. 2A). GAM Model 1 further showed that overall mean cannibalism probability was lower in female than male cannibals (Fig. 2A; β*_Male_* = 1.01, SE = 0.288, t-value = 3.52, p-value < 0.001).

Due to this sex effect on cannibalistic voracity, the upper and lower limits of the predation window were sex-dependent (Fig. 2B). Specifically, the predation window was slightly wider in male (δ = 0.09, ϕ = 0.26, ε = 0.41) than in female cannibals (δ = 0.13, ϕ = 0.26, ε = 0.38). These parameters show that newly-hatched medaka larvae are under strong cannibalistic risk (Fig. 3). An average-sized hatchling (3.8 mm Sdl) may be cannibalised by any 9.3-to-42.2 mm Sdl male (10.0 to 29.2 mm female) conspecific. This body-size range encompasses not only the whole range of parental body sizes, but also late-juvenile body sizes (Fig. 3), indicating that medaka larvae are exposed to both inter- and intra-cohort cannibalism.

**Fig. 3.**
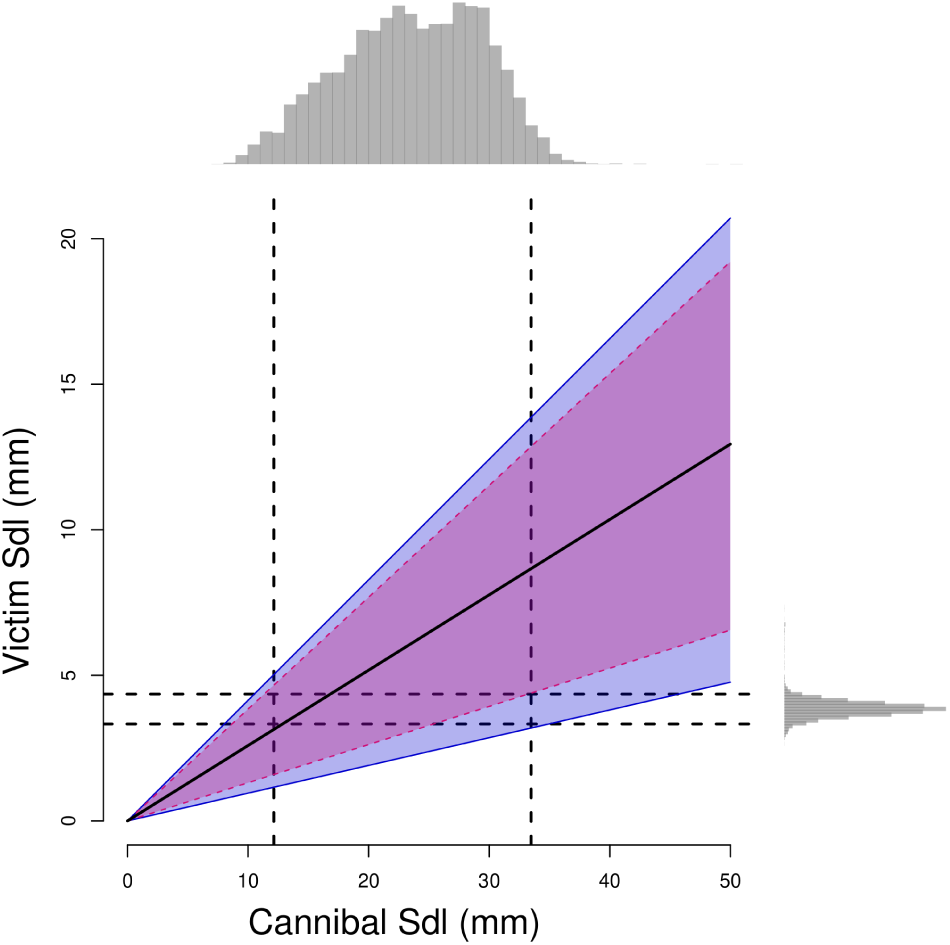
Predation window for victim *vs*. cannibal standard body length (Sdl) in medaka. The black solid line shows the optimal victim length while ribbons show the upper and lower predation limits for males (blue solid lines) and females (pink dashed lines), as defined by the 5 % cannibalism probabilities displayed in Fig. 2B. The top marginal distribution shows the body sizes typically present in pond medaka populations in France before reproduction in March (Bouffet-Halle et al. 2021), i.e., the size distribution of potential cannibals. The right-hand marginal distribution shows larval body sizes at hatch (Renneville et al. 2020), i.e., the size distribution of potential victims. Vertical and horizontal dashed lines show the 2.5 and 97.5 % quantiles of Sdl distributions.

### Pond populations

In replicated pond populations, complex habitats increased the population growth rate (Table 1-Model 2, positive β effect), as indicated by green lines being higher than red lines in Fig. 4A. Specifically, population growth rate increased on average from 1.5 year^-1^ under a low complexity to 2.7 year^-1^ under a high complexity (80 % increase).

**Table 1.**
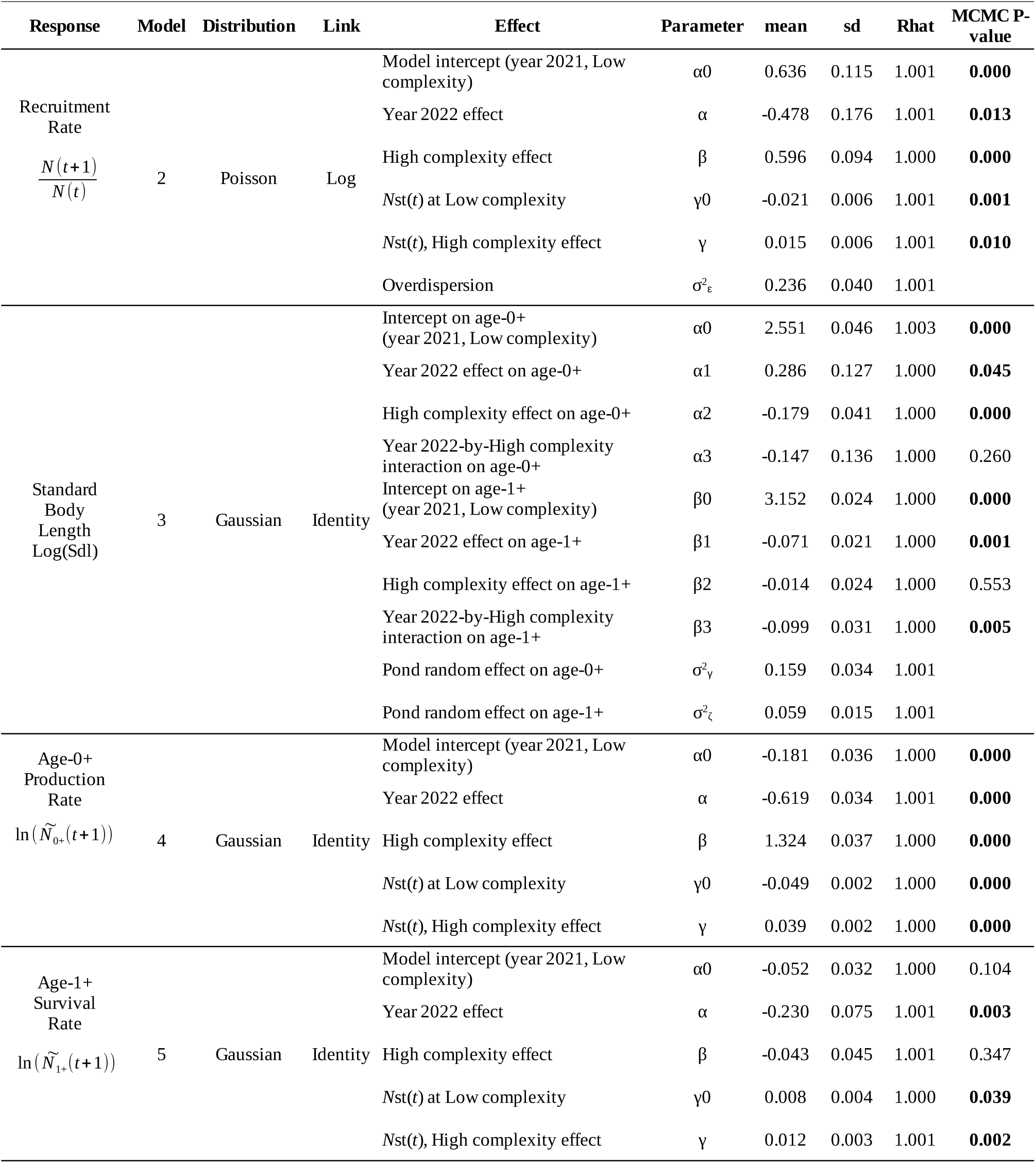
Parameter estimates from Bayesian Models 2-5. The models were fitted using an “effect” parametrization, i.e., intercepts and slopes are evaluated at a reference level (low complexity and year-of-experiment 2021), coded with a “0” in the “Parameter column”, and effects of high habitat complexity and year-of experiment 2022 are estimated as deviations from this reference level. MCMC P-values are twice the proportion of the posterior which sign was opposite to the sign of the posterior mode. P-values significant at the 5 % risk are bold-faced. MCMC P-values are not relevant, and thus not supplied, for variance parameters.

**Fig. 4.**
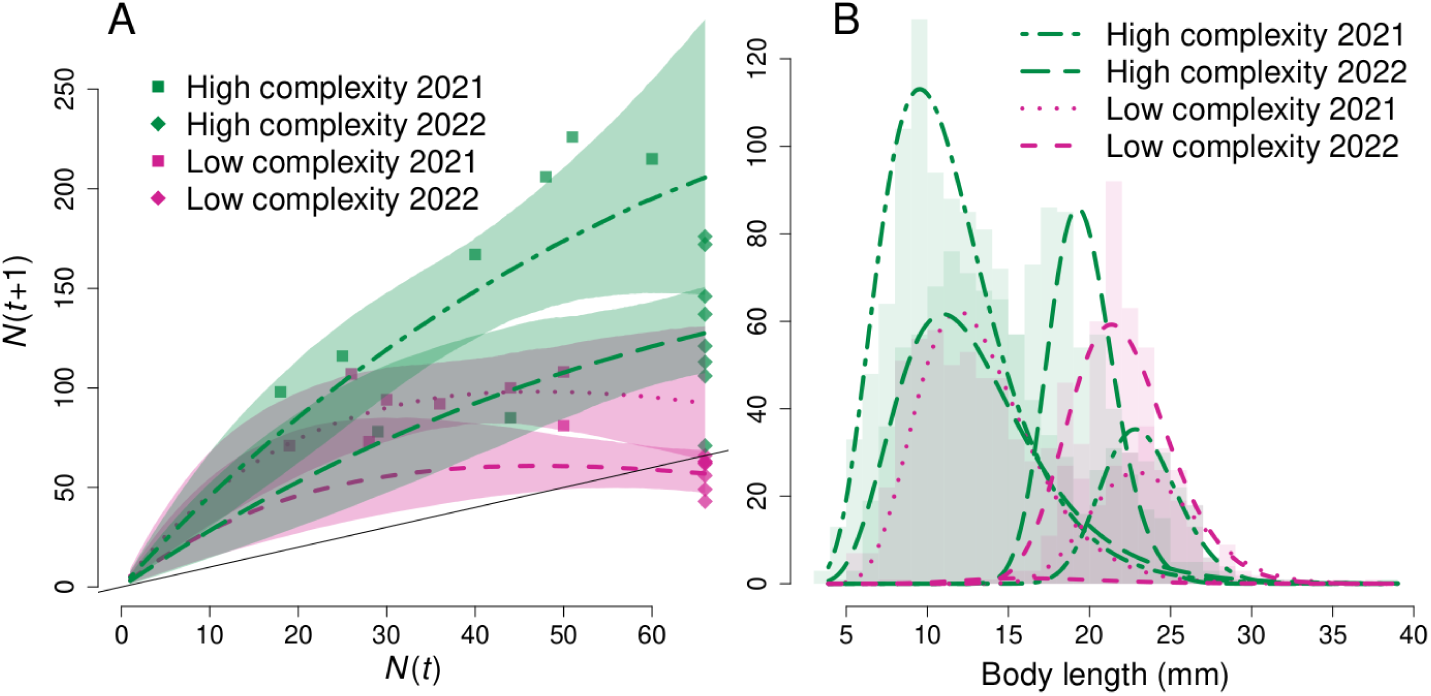
Effect of habitat structural complexity on medaka population dynamics. **A.** Relationship between the number of medaka after (*N*(*t*+1)) and before reproduction (*N*(*t*)) in replicated pond populations. Dots show the raw data (squares: low complexity; diamonds: high complexity), and curves show median posterior predictions from Model 2 (violet-red: low-complexity habitats; Green: high-complexity habitats). Ribbons show 95 % credible intervals. The solid, straight y = x black line intersects coloured curves at habitat carrying capacity K. **B.** Body-length distributions after reproduction at *t*+1. Light bars show the raw data and curves show median posterior distributions predicted from mixture Model 3. There are 8 curves, one per age-class per complexity treatment per year of experiment.

Additionally, medaka population dynamics were negatively density-dependent under a low habitat complexity (Table 1-Model 2, significantly negative γ_0_ effect), but a high habitat complexity dampened this negative density dependence (Table 1-Model 2, positive γ effect). Accordingly, the stock-recruitment relationship changed from flat at low habitat complexity (red lines) to positive at high habitat complexity (green lines), reflecting increased habitat carrying capacity (Fig. 4A). Fitting Model 2 without centring N(t), so as to preserve the original biological interpretation of r (see Methods), yielded a median posterior for K that increased from 82 under low habitat complexity to 194 under high habitat complexity in 2021, and from 59 to 143 in 2022.

Mixture Model 3 allowed us to quantify the effects of habitat structural complexity on age-specific, natural log-transformed body-length distributions. Increased habitat complexity decreased mean age-0+ ln(Sdl) in both years of experiment (Table 1-Model 3, α_2_ and α_3_ parameters), and decreased mean age-1+ ln(Sdl) in 2022, when the complexity contrast was more extreme, but not significantly so in 2021 (Table 1-Model 3, β_3_ *vs*. β_2_ parameters).

In 2021, more complex habitats increased variability in age-0+ ln(Sdl) (Model 3, σ_age-0+, Low-complexity, 2021_^=0.269 *vs*. σ^_age-0+, High-complexity, 2021_^=0.303, MCMC P-value = 0.022), but^ decreased variability in age-1+ ln(Sdl) (Model 3, σ_age-1+, Low-complexity, 2021_=0.140 *vs*. σ_age-1+, High-complexity, 2021_=0.113, MCMC P-value = 0.038). In 2022, age-0+ fish were too few in low-complexity ponds to provide any robust inference on Sdl variability (MCMC P-value = 0.652), while more complex habitats increased age-1+ ln(Sdl) (Model 3, σ_age-1+, Low-complexity, 2022_=0.102 *vs*. ^σ^_age-1+, High-complexity, 2022_^=0.136, MCMC P-value = 0).^

Model 2 is a model of unstructured population dynamics, where age-0+ and age-1+ individuals are lumped in *N* (*t* +1). Hence, from Model 2, it is not possible to tell whether the increase in *N* (*t* +1)/ *N* (*t*) under a high habitat complexity reflected increased *per capita* production rate of age-0+ recruits and/or increased survival probability of age-1+ fish. To answer this question, mixture Model 3 provided us with separate estimates for the numbers of age-0+ and age-1+ fish at *t*+1, and thus allowed us to separate these two effects using Models 4 and and 5 (see Methods).

Model 4, which was fitted to the posterior number of age-0+ medaka, perfectly parallel results from Model 2 above. Specifically, *per capita* production of age-0+ fish strongly increased from on average 0.8 at low complexity to 3.1 under a high habitat complexity (Table 1-Model 4, positive β effect). Additionally, the habitat complexity-by-density interaction reveals a strong negative density dependence under a low habitat complexity (Table 1-Model 4, negative γ_0_ effect) that was dampened under a high habitat complexity (Table 1-Model 4, positive γ effect).

Patterns of age-1+ survival probability, as inferred from Model 5, were opposite to patterns of *per capita* production rates of age-0+ recruits, suggesting opposite effects of habitat complexity on age-0+ and age-1+ fish. Specifically, a high habitat complexity *decreased* age-1+ survival probability in both the Gaussian and binomial models, but significantly-so in the binomial model only (Table 1-Model 5 *vs*. Appendix 3: negative β effects). The binomial model predicted that age-1+ survival probability decreased from 0.91 at low complexity to 0.77 under a high habitat complexity.

Surprisingly, survival probability of age-1+ fish was *positively* density-dependent under low habitat complexity in both the Gaussian and binomial models (Table 1-Model 5 and Appendix 3: positive γ_0_ effects), and a high habitat complexity amplified this positive density dependence, but significantly-so in the Gaussian model only (Table 1-Model 5 *vs*. Appendix 3: positive γ effects), suggesting a weak amplification.

Taken together, results from Models 4 and 5 indicate that the positive effect of habitat complexity on medaka population growth rate resulted from increased *per capita* production of age-0+ fish, and was weakly opposed by depressed survival of age-1+fish.

### Filamentous algae

On average, filamentous algae covered 7.1 % of pond surface in high-complexity ponds, and 13.8 % in low-complexity ponds (Fig. 5). Analysis of Model 6 showed that this difference was statistically significant (α_Low_ = 0.728, SE = 0.259, z-value = 2.81, p-value < 0.005).

**Fig. 5.**
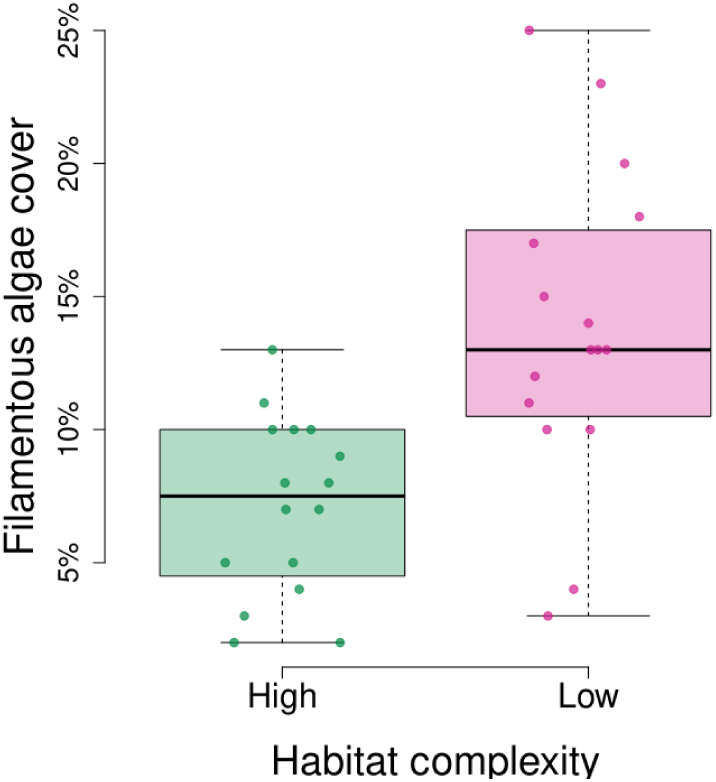
Filamentous algae. Algal cover on each pond was estimated independently by two observers before medaka fishing in November 2022. Symbols show the raw data (n = 32).

## DISCUSSION

In multispecies assemblages, habitat structural complexity increases species coexistence and productivity through a relaxation in the strengths of both interspecific competition and predation (Smith 1972, Crowder and Cooper 1982, Holt 1987, Diehl 1988, 1992, Heck and Crowder 1991, Hixon and Menge 1991, Rogers et al. 2014). Our results show that habitat structural complexity may further increase age-class coexistence, population growth rate and habitat carrying capacity at a single-population level. Hence, intraspecific processes may contribute to the positive effects of habitat complexity on biodiversity and productivity at a community level.

Our results may further be used to gain some understanding of the underlying ecological mechanisms that drive positive habitat complexity-productivity relationships. We show that increased age-class coexistence and productivity in medaka populations were mainly achieved through an increased *per capita* production of age-0+ recruits by their age-1+ parents. In theory, such an increase in recruitment may result from increases in both parental fecundity (i.e., increased egg production) and in larval survival to the recruit stage. However, several lines of evidence point to increased larval survival as the only mechanism at play.

Filamentous algae, one of the main food sources for medaka, were less abundant in high-complexity than in low-complexity ponds, probably due to higher fish densities and/or to artificial structures directly impeding algal growth. As a result of this food shortage, both age-0+ and age-1+ medaka had smaller and more variable body sizes in more complex habitats, indicating increased competition for food (Ohlberger et al. 2013). Age-1+ survival probability further decreased in more complex habitats, consistent with negative effects of food competition on fitness in age-1+ medaka, which are dominated by their age-0+ progeny in exploitative competition for food (Edeline et al. 2016). Therefore, fecundity of age-1+ medaka most likely *decreased* under a higher habitat complexity, and a higher recruitment in more complex habitats is most likely explained by increased survival of age-0+ medaka due to relaxed cannibalism.

Interestingly, habitat complexity did not only relax overall cannibalism, but also the density-dependence of cannibalism (Rosenheim and Schreiber 2022). Specifically, habitat complexity relaxed both (i) the *negative* effect of age-1+ density on age-0+ survival, and (ii) the *positive* effect of age-1+ density on their own survival probability, the later being explained by decreasing densities of dominant age-0+ medaka relaxing the competition for food experienced by competitively-inferior age-1+ medaka (Dionne 1985, Claessen et al. 2000). Relaxed density-dependence in cannibalism shows that habitat structural complexity reduced both the rate of dangerous encounters among medaka, as well as the likelihood that encounters result in successful cannibalistic attacks (Rosenheim and Schreiber 2022).

In the wild in Japan, medaka starve to death when they turn age-1+, while their newly-hatched progeny grow very rapidly (Terao 1985, Awaji and Hanyu 1987, Egami et al. 1988). This “semelparous” life history pattern seems to reflect a competitive exclusion of age-1+ parents by their age-0+ progeny (Edeline et al. 2016). In contrast, in experimental ponds in France medaka survive to age 2+ and contribute to two or more reproductive bouts in their lives, i.e., are “iteroparous” (this study, Bouffet-Halle et al. 2021). In multiple taxa, cannibalism allows large-bodied individuals to overturn the competitive superiority of smaller-bodied conspecifics (this study, Dionne 1985, Polis 1988, Claessen et al. 2000, Wise 2006), and increased cannibalism, as resulting from a lower habitat complexity, is probably an important driver of the extended lifespan of age-1+ medaka in experimental ponds.

Accordingly, wild habitats harbour rooted water plants or wood debris that provide refuges to the juveniles across the whole water column down to benthic habitats, which provide substantial amounts of food to medaka when their preferred planktonic prey are scarce (Terao 1985). In contrast, in experimental ponds complexity was restricted to top water layers. Additionally, wild habitats often harbour shallow and warm areas that are particularly sought after by medaka fish (Awaji and Hanyu 1987), but which expose large-bodied medaka to stranding and thus protect small-bodied juveniles against cannibalism. For instance, in trout (*Salmo trutta*) in lakes, shallow inflowing streams protect juveniles from cannibalism, and more inflowing shallow streams result in increased competition for food among trout in lakes (Uszko et al. 2025). Such shallow areas were absent from our experimental ponds.

If the higher complexity of wild habitats explains relaxed cannibalism and the resultant dominance of age-0+ medaka in the wild, then we could expect a hump-shaped habitat complexity-productivity relationship in medaka populations, with a shift along an increasing habitat-complexity gradient from a cannibal-mediated, age-1+ dominance at low complexity to an exploitative competition-mediated, age-0+ dominance at high complexity, resulting in maximal age-class coexistence and population productivity at intermediate complexity levels where the ontogenetic asymmetry in dominance is most relaxed. Maximal diversity and productivity at intermediate levels of habitat structural complexity are indeed observed in complex, multispecies communities (Crowder and Cooper 1982, Rogers et al. 2018), supporting the contention that complexity-diversity and complexity-productivity relationships emerge through similar, size-dependent predation processes operating at both the community and population levels.

Size-dependent cannibalism is a very common form of interaction in the animal kingdom (Fox 1975, Polis 1981, Smith and Reay 1991, Wise 2006), and cannibalism plays a key role in determining life histories and population dynamics in wild populations of annelids (Elliott 2004), insects (Baskauf 2009), crustaceans (Grosholz et al. 2021), acarids (Walde et al. 1992), fish (Claessen et al. 2000, 2002, Persson et al. 2003), or amphibians (Wissinger et al. 2010). We thus expect habitat complexity, through relaxing cannibalism, to enhance age-class coexistence and population growth rate across a broad taxonomic range. Around the globe, habitat structural complexity becomes increasingly threatened by anthropogenic perturbations such as deforestation, eutrophication, bottom trawling and dredging, river channelization, by trophic cascades resulting from top-predator extinction, or by ocean acidification, thus making global change convergent with a general habitat simplification. Ample evidence already demonstrates that habitat complexity supports species diversity, and that anthropogenic habitat simplification may trigger a cascade of species extinctions in multispecies communities (Smith 1972, Crowder and Cooper 1982, Diehl 1988, 1992, Heck and Crowder 1991, Hixon and Menge 1991, Janssen et al. 2007, Kovalenko et al. 2012, Reichstein et al. 2013, Rogers et al. 2014). Our study further suggests that reduced population growth rate in the surviving species may reduce their resistance to further perturbations, and ultimately impair their longer-term persistence.

## Acknowledgement

We are grateful to the NBRP medaka (https://shigen.nig.ac.jp/medaka/) and to Prof. Kiyoshi Naruse (NIBB Okazaki) for providing us with the medaka strains. We thank André De Roos for insightful comments on an earlier manuscript version. Alexandre Kempf, Solène Moulin, Louisiane Perrin and Morgan Verdeil provided invaluable help during behavioural assays. This experiment was approved by the Charles Darwin Ethical Committee (Ce5/2010/041).

## Author contributions

EE designed and contributed to performing experiments, analysed data and wrote the first draft version. EE, YB and DRR performed experiments. All authors contributed to result interpretation and draft improvements.

## Data, script and code availability

All data and codes needed to run the analyses and to reproduce figures may be downloaded from Zenodo: https://doi.org/10.5281/zenodo.14044171

### Appendix 1. Artificial habitat structure

#### Geotextile 2D structure

To gain some insights about geotextile’s 2D structure, we performed an ImageJ analysis from a top view of a 27 x 42 cm geotextile sample. We found that holes in the mesh represented 38 % of the whole geotextile surface (Fig. S1A). We plotted hole-size distribution, and compared it with the distributions of medaka body width and height, which determine medaka ability to pass through geotextile mesh (Fig. 1B).

The distributions of medaka body width and height were fully included in hole-size distribution (Fig. S1B), indicating that a floating single-layered geotextile tile could not provide any absolute refuge against cannibalism to small-bodied medaka. However, small-bodied medaka had many more holes available to pass through the geotextile than large-bodied medaka, and we hypothesized that the geotextile should selectively hamper the movements of large-bodied medaka.

**Fig. S1.**
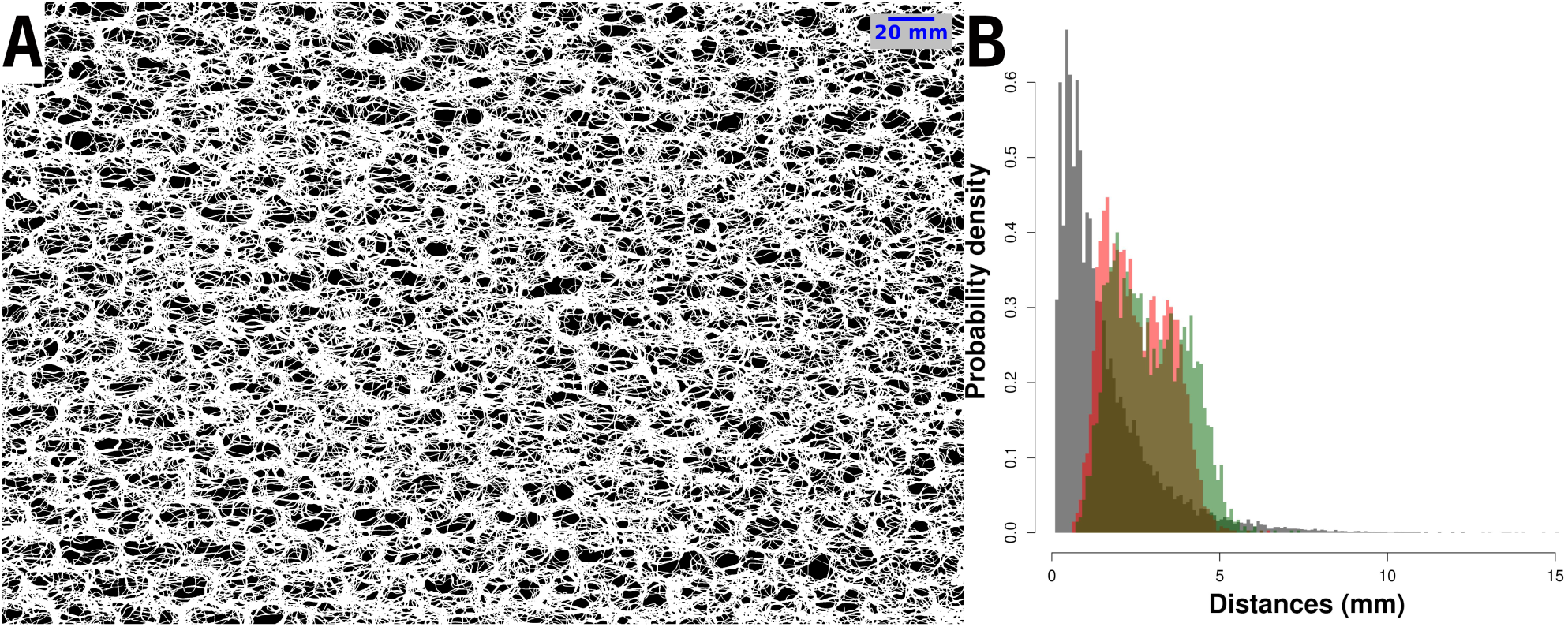
2D structure of a 27 x 42 cm sample of a single layer of geotextile. **A.** Top view of the mesh structure of the geotextile, with holes appearing in black and the geotextile mesh appearing in white. **B. Grey distribution:** distribution of hole size (Feret’s diameter) in the geotextile mesh, as measured using ImageJ from Fig. S1A. Feret’s diameter, also known as maximum caliper, is the longest distance between any two points along the hole boundary. Hole Feret’s diameters ranged from 0.11 to 15.00 mm (mean ± SD = 1.54 ± 1.42 mm). **Red and green distributions:** distributions of medaka body width (red) and height (green) during the 2021 and 2022 mesocosm experiments, as predicted from standard body lengths (Sdl) using allometric relationships estimated by Renneville et al. (2016).

#### Behavioural assays

To test this hypothesis, we evaluated the effect of the geotextile on medaka movements in a series of four behavioural assays performed on July 1^st^ 2024. We used a 55 x 67 cm, single-layered tile of geotextile affixed onto a wooden frame and placed in a 100 x 100 cm tank filled with 30 cm of dechlorined tap water (Fig. S2). The frame was put on wedges, such that the geotextile was maintained 3-4 cm below the water surface, and such that the fish put inside the frame would have to swim through the geotextile to escape to the tank.

**Fig. S2.**
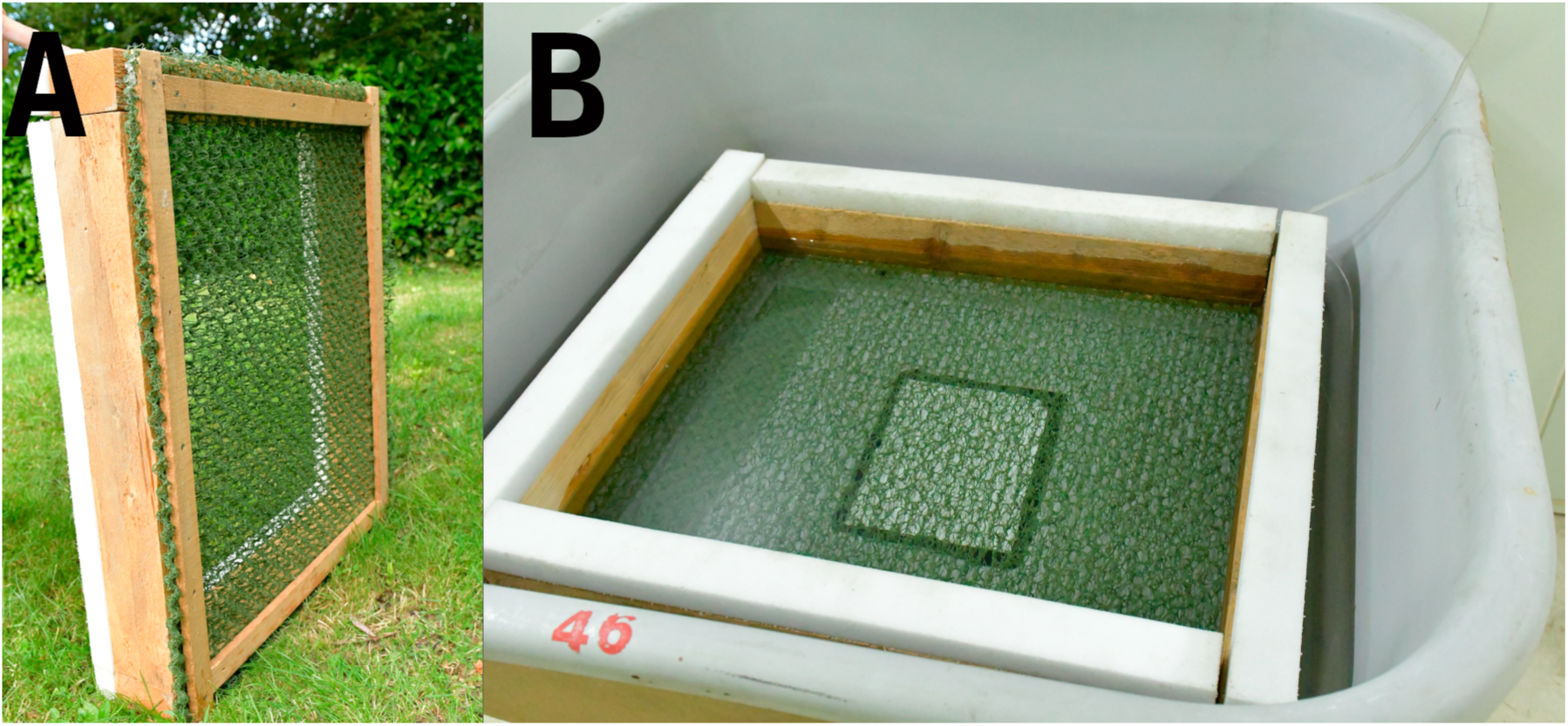
Single-layered geotextile tile affixed onto a wooden frame. **A**: Bottom view of the 55 x 67 cm wooden frame. **B**. Top view of the wooden frame placed in the tank before the start of behavioural assays.

The medaka used for assays were from a stock of the Kiyosu strain maintained permanently in large outdoor mesocosms with mild artificial feeding. Four batches of 210, 256, 251, and 220 naive fish, respectively, were gently captured using an aquarium net. With no further habituation, each batch of fish was deposited in the wooden frame and left undisturbed under room light during 1, 5, 10 or 20 minutes, respectively. At the end of each assay, the frame was removed from the tank and fish were separated among those that had swum through the geotextile and those that did not. Each individual fish was assayed only once. All fish were photographed and measured for standard body length (Sdl) using ImageJ as described in the main text, and were returned to their outdoor mesocosm.

We modelled the effect of medaka Sdl (range 8-32 mm, mean ± SD = 20.4 ± 4.8 mm) and duration of assay on medaka probability to swim through the geotextile using a Bernouilli GLM:

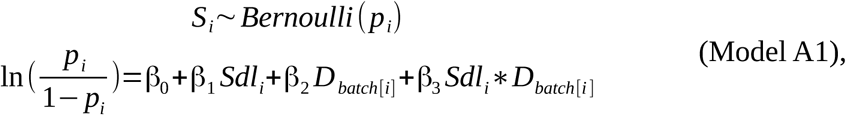

where *S_i_* is the status of individual fish *i* at the end of the behavioural assay, i.e., *S_i_*=1 if the fish swam through the geotextile and 0 otherwise, and *D_batch_*_[_*_i_* _]_ is the duration of assay associated with the batch to which fish *i* belonged. The results of maximum-likelihood estimation of the β parameters using the glm function of R are shown in Table A1.

**Table A1.**
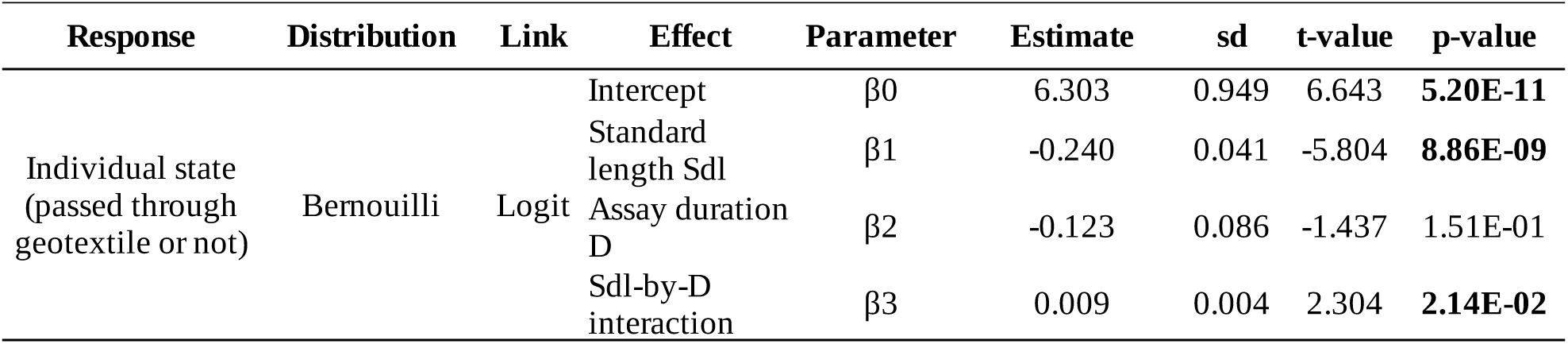
Results of maximum-likelihood estimation of parameters from model A1. Bolded-faced p-values are significant at the 5 % risk.

| Response | Distribution | Link | Effect | Parameter | Estimate | sd | t-value | p-value |
| --- | --- | --- | --- | --- | --- | --- | --- | --- |
| Individual state<br>(passed through<br>geotextile or not) | Bernouilli | Logit | Intercept | $\beta_0$ | 6.303 | 0.949 | 6.643 | **5.20E-11** |
| | | | Standard length Sdl | $\beta_1$ | -0.240 | 0.041 | -5.804 | **8.86E-09** |
| | | | Assay duration D | $\beta_2$ | -0.123 | 0.086 | -1.437 | 1.51E-01 |
| | | | Sdl-by-D interaction | $\beta_3$ | 0.009 | 0.004 | 2.304 | **2.14E-02** |

Parameter estimation shows that the probability for a medaka to swim through the single-layered geotextile tile was negatively body-length dependent, and that there was a significant length-by duration interaction (Table A1). To further visualize this interaction, we plotted model predictions in Fig. S3.

**Fig. S3.**
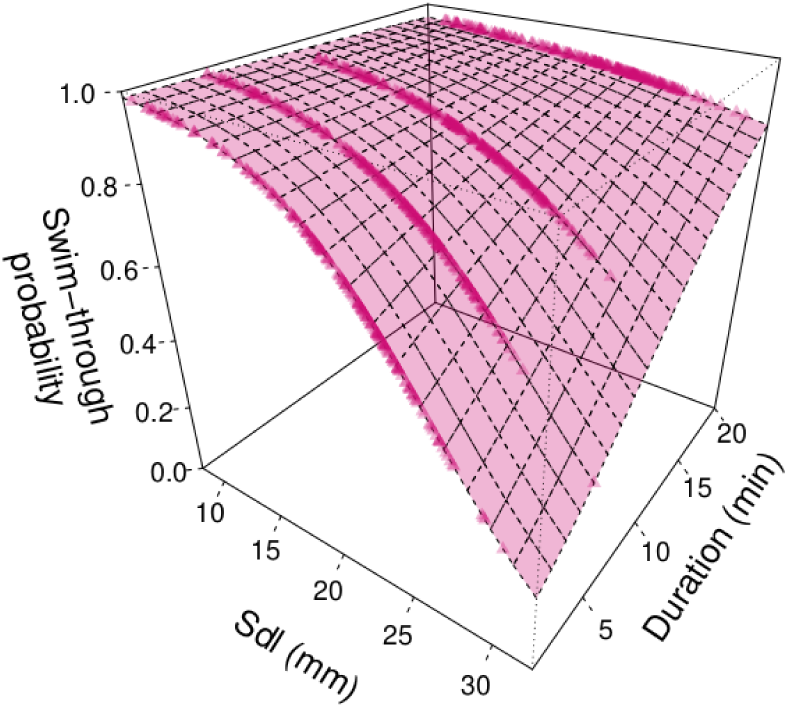
Interaction between medaka standard body length (Sdl) and behavioural-assay duration on medka probability to swim through a single layer of geotextile. The surface shows predictions from Model A1 for the whole range of assayed body sizes. Symbols show the location of fitted values.

Fig. S3 shows that the geotextile strongly reduced passage of the largest-bodied medaka during short-duration assays (1-5 minutes), but almost not during the 20-min assay. This result confirms that a single-layered geotextile tile slowed down, but did not prevent, the movements of large-bodied, potentially cannibalistic medaka.

#### Multiple-layered geotextile mats

During the 2021 pond experiment, we used single-layered geotextile tiles and varied habitat complexity through contrasted tile surfaces. In 2022, we aimed at increasing the complexity contrast, and we varied complexity through contrasting both geotextile surface and the number of geotextile layers. Specifically, we created high-complexity habitats using geotextile mats comprised of five stacked tiles, which structure is shown in Fig. S4.

**Fig. S4.**
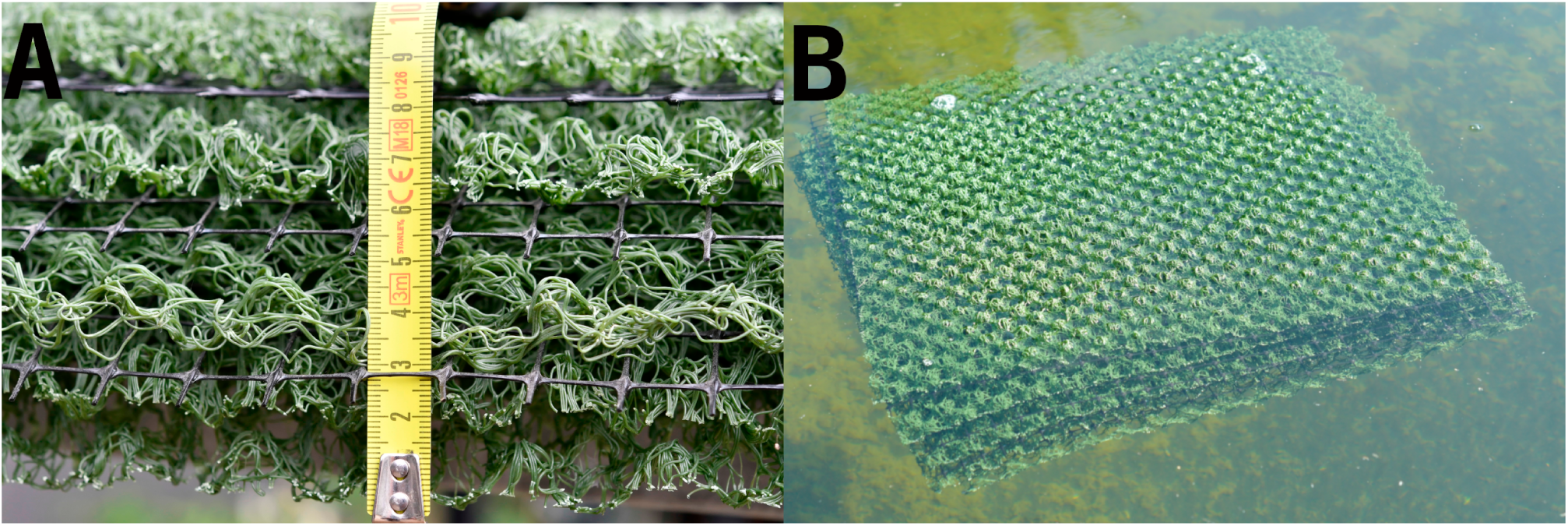
Geotextile mat made from a 5-layered stack of geotextile tiles. **A.** Close view showing the 5 layers of geotextile, separated by a 2 cm x 1 cm black plastic mesh used to prevent tiles to interlock. This mesh was large enough not to restrict fish movements. The whole mat was about 10 cm thick. **B.** Distant view showing mat positioning at the top of the water column. Mat layers were bound together by a pair of plastic collars, here visible on the right-hand side of the mat.

### Appendix 2. Strain effects on population growth rate and body sizes

In the 2021 pond experiment, medaka strains were not mixed in a pond (see Methods), and it was possible to test for strain effects on population dynamics. We tested for a strain effect on population growth rate using a model similar to Model 2 in the main text:

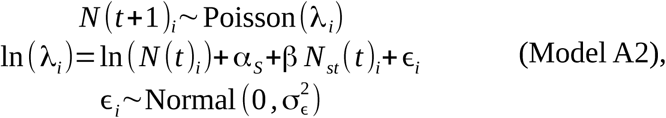

where *i* indexes ponds (n = 16), α*_S_* is a strain effect on population growth rate, β is the density-dependent parameter, and σ^2^ is a overdispersion parameter. Model A2 was fitted in the glmmTMB library of R (Brooks et al. 2017). We tested for a stain effect using a Chi-squared likelihood ratio test between Model A2 and a similar model that did not include any strain effect using the anova function of R. The results suggest no strain effect on population growth rate (Table S2).

**Table S2.** Results from running the anova function in R to compare Model A2 with a similar model that omits a strain effect.

|  | Df | AIC | BIC | logLik | deviance | Chisq | Chi Df | p-value |
| --- | --- | --- | --- | --- | --- | --- | --- | --- |
| Model without any strain effect | 3 | 165.8 | 168.2 | -79.9 | 159.8 |  |  |  |
| Model A2 | 8 | 167.6 | 173.8 | -75.8 | 151.6 | 8.2 | 5 | 0.144 |

We tested for a strain effect on standard body lengths Sdl using a linear mixed-effects model fitted by restricted maximum likelihood in the nlme library of R (Pinheiro et al. 2018):

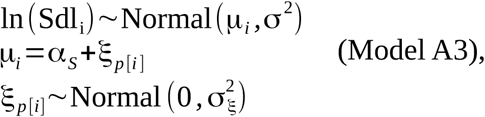

where ln is the natural logarithm, *i* indexes individual fish (*n* = 1917), *p* indexes ponds (*n* = 16), and α_S_ captured strain effects on mean ln-transformed body lengths. The ξ*_p_* [*_i_*] parameter captured random pond effects. We then tested significance of the variance explained by α_S_ in the model using an F-test in the anova function of R. The results show that the strain effect was not statistically significant (Table S3).

**Table S3.** Result of running the anova function in R to test significance of a strain effect in Model A3.

|  | **numDF** | **denDF** | **F-value** | **p-value** |
| --- | --- | --- | --- | --- |
| (Intercept) | 1 | 1901 | 5122.4 | <.0001 |
| Strain | 5 | 10 | 1.0 | 0.451 |

### Appendix 3. Binomial analysis of survival probability of age-1+ medaka through the reproductive period

Compared to Model 5 in the main text, an alternative approach to modelling survival probability of age-1+ medaka, which does not propagate estimation uncertainty in age-1+ numbers but preserves the underlying Binomial process, consists in using an overdispersed Binomial model of the form:

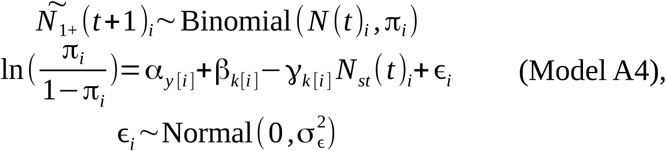

Where 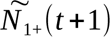 is the median posterior number of age-1+ fish after their reproductive period, and ɛ is an overdispersion parameter. Other variables and parameters are as in Model 5 in the main text. Model A4 had a Bayesian P-value of 0.49, indicating an excellent fit to the data, and yielded parameter estimates presented in Table S4.

**Table S4.** Summary statistics for MCMC estimation of the parameters of Model A4. Bolded-faced p-values are significant at the 5 % risk.

| Response | Distribution | Link | Effect | Parameter | mean | sd | Rhat | MCMC P-value |
| --- | --- | --- | --- | --- | --- | --- | --- | --- |
| Age-1+ Survival probability $\pi$ | Binomial | Logit | Model intercept (year 2021, Low complexity) | $\alpha_0$ | 2.386 | 0.657 | 1.037 | **0.000** |
| | | | Year 2022 effect | $\alpha$ | -1.747 | 1.067 | 1.022 | 0.132 |
| | | | High complexity effect | $\beta$ | -1.154 | 0.583 | 1.017 | **0.047** |
| | | | $N_{st}(t)$ at Low complexity | $\gamma_0$ | 0.098 | 0.039 | 1.046 | **0.026** |
| | | | $N_{st}(t)$, High complexity effect | $\gamma$ | 0.002 | 0.034 | 1.025 | 0.946 |
| | | | Overdispersion | $\sigma_\epsilon^2$ | 1.527 | 0.249 | 1.005 | |

## Footnotes

1 https://shigen.nig.ac.jp/medaka/

